# Semi-supervised Retrieval of Functional Residues Through the Integration of Protein Language Models and Gene Ontology Data

**DOI:** 10.64898/2025.11.28.691208

**Authors:** Andrew Dickson, Safaa Mouline, Ali Tamadon, Mohammad R.K. Mofrad

## Abstract

**Motivation:** Experimental studies of protein function often focus on mechanistic descriptions, characterizing how specific sites and residues contribute to activity. Abstractions such as domains and active sites enable quantitative descriptions of how protein features act biologically. Thanks to the abundance of high-quality sequence and function data, machine learning has achieved great success in directly predicting protein function. However, translating functional characterizations into mechanistic ones on the level of the domains, binding sites, or motifs remains challenging. This represents a semi-supervised problem: sequences and global functional labels are available, but local annotations must be inferred.

**Results:** We investigate the unsupervised discovery of functionally active protein regions by integrating protein sequence models with functional information. We first formalize the residue-level functional annotation problem by constructing unified evaluation datasets linking Gene Ontology functions to annotated residues. Eight datasets are assembled, spanning levels of specificity from single active-site residues to domains covering up to 60% of a protein. We then introduce a new class of function-conditioned generative models that more accurately predict functionally important residues than existing approaches, including interpretability methods and PSSM entropy estimation, across multiple benchmark datasets.

**Availability:** github.com/mofradlab/go_interp

## 1. Introduction

### 1.1. Problem and Gap

While protein machine learning models achieve high performance on functional classification tasks Boadu, A Lee, and Cheng 2025, it is difficult to directly characterize the elements of a protein responsible for their function. Often experimental data comes in the form of high level information on function, such as knock-out experiments, or the results of high-throughput experiments, in which sequence level information on protein function is available, but residue level annotations may be far more difficult to acquire.

It is common to build databases of conserved regions, motifs, or sites, and functionally annotate them through comparison to real world experiment, such as in the Conserved Domain Database (CDD) Marchler-Bauer et al. 2011, or the PRINTS database of protein fingerprints Attwood et al. 2003.

### 1.2. Prior Methods and Limitations

While there are dozens of existing methods for predicting conserved elements in proteins, most rely purely on sequence or evolutionary information. Prior methods typically rely on global importance estimates, in which the general relevance of residues are estimated through methods such as sequence conservation, and these are used for a proxy for functional importance. Sequence-based methods, such as ConSurf Ashkenazy, Abadi, et al. 2016, SIFT Sim et al. 2012, and those using information-theoretic measures like Jensen-Shannon Divergence (JSD) Capra and Singh 2007, typically estimate a position-specific scoring matrix (PSSM) for residues using multiple sequence alignments of homologous sequences. Common pipelines include a homology search, such as through PSI-BLAST, followed by multiple sequence alignment through tools such as MAFFT or MUSCLE Altschul et al. 1997, Katoh and Standley 2013, Edgar 2004. At the same time, modern pre-trained generative models over protein sequences, like those from the ESM (Evolutionary Scale Modeling) family Lin et al. 2023, may be used to estimate the same scoring statistics directly without needing an explicit alignment Meier et al. 2021.

However, because these methods ignore experimental functional information they cannot distinguish conservation for functional reasons from conservation for reasons such as structural stability or alternative functions Pancsa and Fuxreiter 2012. Annotated datasets, such as the CDD or PRINTS, or larger aggregates such as InterPro Hunter et al. 2008, typically rely on post-hoc annotation of identified regions, instead of directly integrating functional information into sequence annotations. There is a need for methods capable of accounting for prior knowledge of protein function, but the effectiveness, and feasibility, of potential approaches are unclear.

### 1.3. Contributions

In this work, we formalize semi-supervised functional residue prediction by constructing a benchmark of eight datasets. We then systematically compare the retrieval performance of conservation-based methods, classifier interpretability techniques, and generative language models. Finally, we introduce a novel function-conditioned generative model trained with a context-span masking objective for the purpose of incentivizing the use of functional information.

A natural way to predict functional information is to extract residue level predictions from function classifier models, such as predictors of Gene Ontology (GO) annotations Ashburner et al. 2000, Camon et al. 2004. Existing machine learning models trained on annotations already outperform sequence search based methods Sanderson et al. 2023, and in particular, pre-trained protein language models (pLMs) fine-tuned on annotation data show great success in producing high-quality annotations Littmann et al. 2021. We apply multiple interpretability algorithms to a strong pLM for a residue retrieval task. At the same time, we compare against several sequence conservation based approaches using MSAs or generative models.

We find that in most cases a generative modeling, entropy-based, approach is stronger than extracting residue-level information from even a strong classifier. Naive training for classification appears to lose or ignore information relevant to the sequence distribution, losing important residue level information. Notably, we do find that in cases of high-entropy, high-relevance residues such as those in the Eukaryotic Linear Motif resource Kumar et al. 2019, interpretability based methods show the strongest overall performance.

For most problem settings, we find that functional information, in the form of global assignment of Gene Ontology terms to protein sequences, can be integrated into predictions of residue importance through the correctly incentivized fine-tuning of generative sequence models. For the best overall retrieval of function-relevant residues with an unsupervised method, we develop a protocol for fine-tuning pre-trained models to solve an artificially challenging masked language modeling objective conditional on functional information.

## 2. Materials and Methods

### 2.1. Benchmark Datasets

For our evaluations, we build eight datasets with residue-level annotations fig. 1. Two datasets identify active sites from the Catalytic Site Atlas and InterPro, and two identify binding sites from BioLiP2 and InterPro Ribeiro et al. 2018, Hunter et al. 2008, C Zhang et al. 2023. We also include a dataset of short linear motifs from the Eukaryotic Linear Motif resource, a dataset of regions driving liquid-liquid phase separation (LLPS) from PhaSePro, and finally, two datasets of functionally implicated repeats and domains, also extracted from InterPro Kumar et al. 2019, Mészáros et al. 2020. InterPro is an aggregate of annotated regions from over a dozen datasets, which we broadly decompose into four classes of entries: those representing active sites, binding sites, repeat regions, and domains. We then build each of our four datasets by filtering InterPro for those entries involved with SwissProt proteins and downsampling to a small set of representative region identifiers.

**Fig. 1:**
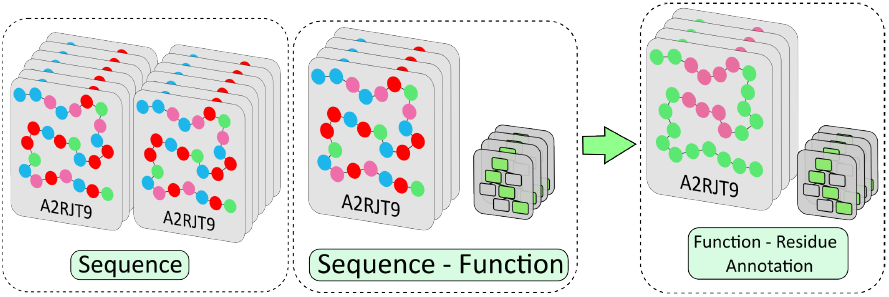
(A) Dataset structure for the residue level functional annotation problem. Protein sequence, and sequence-function datasets may be constructed from large existing databases. From these, we construct and evaluate position-level models designed to recover functionally active residues.

Each dataset is filtered to proteins of length of at most 850 residues containing regions associated with at least one gene ontology term. In the case of the CSA, we convert enzyme commission (EC) identifiers to gene ontology terms International Union of Biochemistry and Molecular Biology. Nomenclature Committee 1992, Ashburner et al. 2000, while for the LLPS datasets, we manually associate plain text descriptors with gene ontology terms.

Because of their large size, and to produce datasets of comparable sizes to the CSA, ELMS, and LLPS datasets, we downsample the BioLip and InterPro derived datasets to between 100 and 1000 proteins table 1. In the case of BioLiP, we select out the top 12 most common binding ligands and sample 50 proteins for each. For each selected InterPro category, we sample 12 representative identifiers, and for each identifier sample 50 associated proteins. We then filter out domains or regions covering over 60% of a protein sequence, and annotate regions using the interpro2go database Hunter et al. 2008.

**Table 1.** Summary of Dataset Statistics.

| Dataset | Residues | Annotations | Perc. Coverage |
| --- | --- | --- | --- |
| CSA | $3.00 \times 10^5$ | $3.59 \times 10^3$ | 0.01 |
| IP Active Site | $1.67 \times 10^5$ | $5.56 \times 10^3$ | 0.04 |
| IP Binding Site | $1.87 \times 10^5$ | $8.51 \times 10^3$ | 0.06 |
| ELMS | $1.06 \times 10^5$ | $2.16 \times 10^3$ | 0.03 |
| BioLip | $2.50 \times 10^5$ | $5.51 \times 10^3$ | 0.04 |
| LLPS | $2.98 \times 10^4$ | $1.10 \times 10^4$ | 0.37 |
| IP Repeat | $1.45 \times 10^5$ | $1.73 \times 10^4$ | 0.15 |
| IP Domain | $8.76 \times 10^4$ | $2.96 \times 10^4$ | 0.33 |

**Table 2.** Comparison of PSSM and demasking entropy for annotation retrieval. Performance was evaluated across four datasets using Area Under the ROC Curve (AUC), Mean Reciprocal Rank (MRR), and recall for the top 30 predictions (Top-30). ESM-C demasking outperforms PSSM across almost all metrics and datasets.

| Dataset | MSA |  |  | ESM-C |  |  |
| --- | --- | --- | --- | --- | --- | --- |
|  | AUC | MRR | Top-30 | AUC | MRR | Top-30 |
| CSA | 0.79 | 0.19 | 0.42 | 0.82 | 0.40 | 0.59 |
| ELMS | 0.50 | 0.05 | 0.09 | 0.52 | 0.077 | 0.09 |
| LLPS | 0.38 | 0.30 | 0.24 | 0.45 | 0.33 | 0.30 |
| IP Domain | 0.74 | 0.62 | 0.61 | 0.79 | 0.79 | 0.70 |

### 2.2. Sequence Resources and GO Annotations

We construct a dataset of high quality protein sequences and functional annotations using the SwissProt Camon et al. 2004, Gene Ontology Annotation database, and Gene Ontology datasets using our previously developed gobench.org web application Dickson et al. 2023. We filter for annotations sourced from human-reviewed experimental codes International Union of Biochemistry and Molecular Biology. Nomenclature Committee 1992 selected and apply otherwise default settings. In total, the dataset contains 112,000 proteins, with an average of ten annotations for each protein. We apply an 85-15 train-validation split for hyperparameter tuning, but do not benchmark GO annotation models outside of downstream performance on interpretability tasks.

### 2.3. PSSM Construction

To construct Position-Specific Scoring Matrices (PSSMs), we first searched the UniRef90 database Suzek et al. 2015 using MMSeqs2 Steinegger and Söding 2017. After filtering homologous sequences based on identity and length, we generated Multiple Sequence Alignments (MSAs) with Muscle5 Edgar 2022. From the MSA, PSSM scores were calculated from amino acid counts at each position, with residue columns having ¿30% gaps assigned maximum entropy, following Capra and Singh 2007.

### 2.4. Protein Language Models and Training

For classifier interpretability and generative modeling methods for predicting functional importance, we use the pre-trained protein language model, ESM-Cambrian (ESM-C). For function prediction, we fine-tune the model on Gene Ontology (GO) term classification, achieving a maximum F1-score of 0.83 on our held-out dataset. For generative modeling, we introduce a method to condition the ESM-C model on GO terms to generate protein sequences fig. 2. We find that a novel context-span masking objective is necessary for the model to most exploit functional information during generation.

**Fig. 2:**
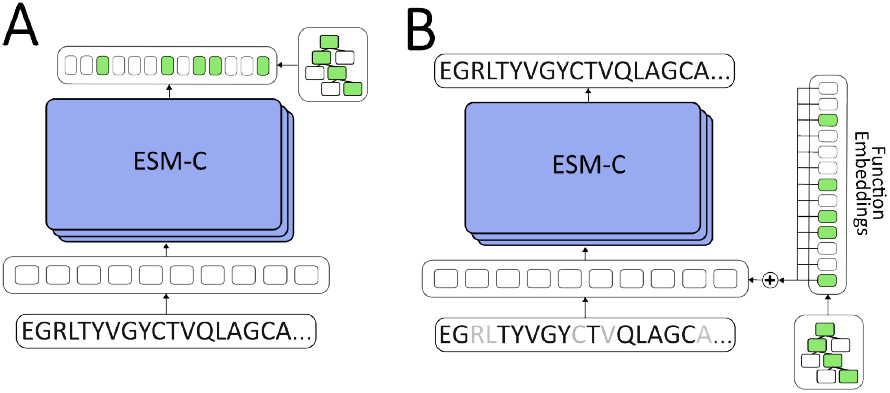
A) The fine-tuning architecture for a GO classifier model. We replace the token logit output of the pre-trained sequence model with a biased linear transform of the average of all residue embeddings from the final layer of ESM-C. Outputs are treated as logits for each Gene Ontology class, and trained with binary cross entropy. B) The fine-tuning architecture for a function conditioned masked modeling model over protein sequences. While the pre-trained ESM-C output is left unchanged, the input residue sequences are perturbed by a sequence level functional embedding.

Our work uses a dataset 𝒟 of *N* protein sequences, 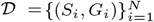. Each protein sequence *S*_*i*_ is a series of amino acid residues, *S*_*i*_ = (*r*_1_, *r*_2_, … , *r*_*L*_), where *L* is the sequence length. The function of each protein is described by a set of Gene Ontology (GO) terms *G*_*i*_ ⊆ 𝒢, where 𝒢 is the complete set of GO terms. For classification, we represent *G*_*i*_ as a multi-hot binary vector *y*_*i*_ ∈ {0, 1}^|𝒢|^.

The primary model used for fine-tuning and generative modeling was the 600M ESM-Cambrian protein language model, trained on a masked language modeling objective derived from the original BERT training procedure Lin et al. 2023, Devlin et al. 2019, Hayes et al. 2024. For managing sequence data, we use the BioPython package Cock et al. 2009. For machine learning training and inference we use the PyTorch and PyTorch Lightning packages Paszke et al. 2019, Falcon and The PyTorch Lightning team 2019. For evaluation we use the Pandas, NumPy, and Scikit-learn packages team 2020, Harris et al. 2020, Pedregosa et al. 2011.

We adopt a two-phase fine-tuning strategy for our GO classification problem setting. Let the ESM-C encoder be denoted by *f*_*θ*_, where *θ* represents the model parameters. For a given sequence *S*, the model produces final-layer residue embeddings *H* = (*h*_1_, *h*_2_, … , *h*_*L*_) = *f*_*θ*_(*S*), where *h*_*i*_ ∈ ℝ^*d*^.

These embeddings are averaged to produce a sequence-level representation, which is passed to a linear classification head. The final probabilities *ŷ* are obtained by applying an element-wise sigmoid function, *ŷ* = *σ*(*z*) fig. 2.

The classification model is trained by minimizing the binary cross-entropy (BCE) loss over all GO annotations:

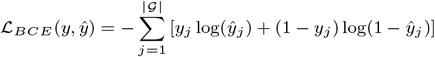

The first fine-tuning phase trains only the classification head (*W, b*) with the ESM-C parameters *θ* frozen, using a high learning rate. The second phase fine-tunes the entire model (*θ, W, b*) for five epochs with a low learning rate annealed from 1×10^−5^ to 1×10^−6^. Our final classifier achieves a maximum F1-score of 0.83 on our validation data, averaged over all GO classes.

### 2.5. Function Conditioned Fine-tuning

Given our dataset of paired protein sequences and GO terms, we also fine-tune the generative sequence model to condition on functional information. We augment the pre-trained ESM-C model with a learnable GO term embedding matrix 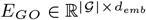.

During evaluation of a protein with associated GO terms *y*, the corresponding embedding vectors are summed to create an aggregate function vector 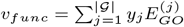.

A linear transform of this aggregate vector is then added to each initial residue embedding *x*_*i*_ at the first layer of the ESM-C transformer to perturb the model’s output: 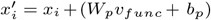 where *W*_*p*_ and *b*_*p*_ are learnable projection parameters. The ESM-C and function embedding parameters are then jointly fine-tuned on a masking loss objective.

We then fine-tune the function-conditioned ESM-C model on the standard masked modeling task, where a set of random indices *M*_*rand*_ corresponding to 15% of the sequence residues are masked. The loss is then.

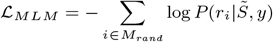

### 2.6. Context Span Masking Objective

During evaluation, we find that function conditioned fine-tuning on a BERT-style random masking objective produces results almost identical to baseline, indicating that functional information is not sufficiently utilized by our model under the default training objective section 3.3. We hypothesized that the standard masked language modeling objective is too simple to force the model to account for global functional information.

Masking contiguous spans is known to increase the difficulty, and potentially the quality, of masked language modeling Joshi et al. 2020, while protein models are known to rely in part on full-sequence context to infer domain distributions Z Zhang et al. 2024. Based on both results, we developed a more challenging training objective, termed context-span masking, designed to obscure larger, contiguous blocks of local information, and remove all knowledge of the protein sequence outside of a small context window.

In this modified task, models are forced to focus on contiguous subsequences. For a sequence of length *L*, we define a random focus region of length one hundred within the full protein sequence. Residues outside this region are masked and excluded from the loss calculation. Within the focus region, residues are masked in contiguous spans of 2 to 5 residues until 15% of the total residues in the sequence are masked, and the model is trained on the same cross-entropy loss.

For both fine-tuning processes, the pre-trained model is fine-tuned with functional conditioning for five epochs at a learning rate of 1 × 10^−5^ annealed to 1 × 10^−6^. As a baseline, we also fine-tune the base ESM-C model on the same masking objectives and dataset without functional conditioning.

### 2.7. Estimating Entropy and Entropy-Gain

PLM predictions are known to estimate the likelihoods of potential residue mutations Meier et al. 2021. To estimate a position specific scoring matrix from a masked language model, we average predictions of the categorical distribution of each residue **esm**, Z Zhang et al. 2024. A large batch of masked sequences is generated from a target sequence such that each residue is masked at least once within the batch. We then estimate the categorical distribution of amino acids at each masked position with application of our model and average over all predicted distributions for each position to produce a PSSM estimate.

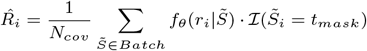

We may then estimate a positional entropy 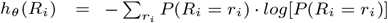.

For the base ESM-C model we randomly mask batch elements until fifteen percent of the total are overwritten, and generate batches of masked sequences large enough that the amino acid distribution for each residue is predicted at least six times in total. Based on calibration tests on the CSA dataset, we find that for pre-trained models retrieval performance improves with the number of masked predictions for each residue, but that performance saturates quickly and is largely robust to masking strategy (see Table S1, S2).

For models fine-tuned on the context-span masking objective, we generate context-span formatted masks for batches, and average only over masked tokens within the open window region, to match the fine-tuning training objective.

To isolate the effect of function conditioning in masked language modeling, we take the additional step of calculating entropy gain scores for our fine-tuned generative models. Here, entropy estimates from function-conditioned models are subtracted from the entropy estimates of baseline models fine-tuned on the same data without access to functional information section 3.3. The resulting scoring function is 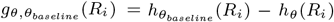, which corresponds to the reduction in entropy of a residue’s distribution from the inclusion of functional information fig. 5.

### 2.8. Interpretability Methods

Given a strong classifier, interpretability methods provide a method for predicting the importance of features to the model’s output, implicitly predicting the importance of residue features to GO functions.

To attribute classifier predictions to specific residues, we employed four methods from the Captum package Kokhlikyan et al. 2020: Layer-wise Integrated Gradients Sundararajan, Taly, and Yan 2017, Gradient X Activation Sundararajan, Taly, and Yan 2017, DeepLIFT Shrikumar, Greenside, and Kundaje 2019, and GradientShap Lundberg and S.-I Lee 2017. For each method, we used the L2 norm of the attribution scores over the residue embedding dimensions to produce a single importance score per residue.

### 2.9. Evaluation Metrics

Scoring methods are evaluated either by the rank they assign functionally annotated residues or by their ability to positively classify functionally annotated residues.

We calculate each rank-based method on a per-protein basis, and then average over proteins for a dataset level statistic. Scores over protein residues are converted to importance rankings.

To evaluate the number of retrievals required to find a first relevant residue, we calculate the mean reciprocal rank (MRR) Radev et al. n.d. For a given protein, the reciprocal rank corresponds to the inverse of the lowest rank of a relevant residue. We average over all proteins to calculate MRR. We apply MRR in particular to the case in which a small percentage of protein residues are relevant, such as for datasets of active or binding sites.

To evaluate the percentage of relevant residues in a small initial set of retrievals, we also calculate the fraction of the top thirty ranked residues which are in the set of functionally relevant positions Capra and Singh 2007. For proteins with less than thirty relevant residues, the top-30 metric is reweighted so that placing all relevant results within the first thirty residues will receive a perfect score of 1.0. Top-30 percentages are averaged over all proteins for the final dataset metric.

For meanAUC calculation, we adopt methods previously applied to sequence conservation estimates Capra and Singh 2007, in which the area under the reciever-opererator characteristic curve (ROC-AUC) is calculated for reach protein, with functionally-annotated residues as positive datapoints and the remainder as negative. AUC scores are then averaged to produce the meanAUC for a dataset.

## 3. Results

### 3.1. Masked modeling entropy estimates outperform MSA

Both multiple sequence alignment and generative masking methods estimate the position specific distribution over residues for a protein. As an initial baseline, we compared their performance across the CSA, ELMS, LLPS, and InterPro domain datasets, selected for their diversity in specificity and function. Both MSA and masked modeling based PSSMs are converted into positional importance values by calculating the entropy of the categorical distribution over possible amino acids.

While both estimates produce similar results on annotation retrieval across datasets, ESM-C demasking predictions outperform MSA across the board, with between a 2 and 5 percent increase in meanAUC performance for most datasets, and a comparable increase across almost every metric **??**. Notably, both methods are worse than random for the case of retrieving important disordered domains, although demasking does achieve considerably higher performance overall. Because of the dominance of masked modeling for residue retrieval across datasets, and the high computational cost of constructing complete MSAs, we restrict our focus to masked modeling for following benchmarks.

### 3.2. Benchmarking of Classifier Interpretability Methods

To determine the relative effectiveness of interpretability methods, we apply the gradient x activation, layer deep lift, integrated gradients, and SHAP methods to each of our assembled datasets of proteins. table 3. Because each method assigns model features importance scores to a single output, we apply each to the classifier prediction of the most specific GO term associated with a target protein.

**Table 3.**
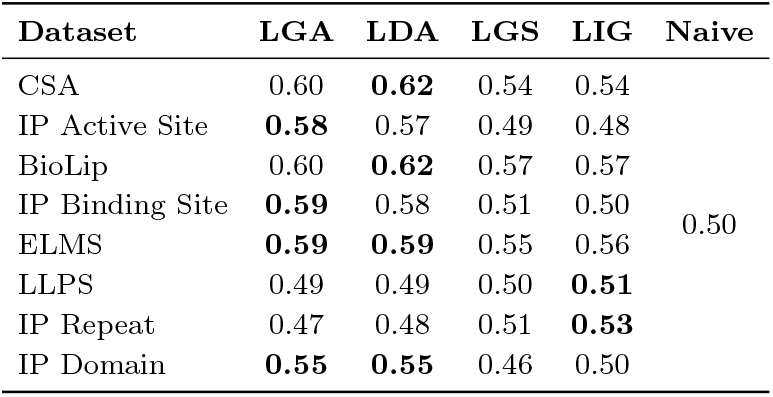
meanAUC scores for each interpretability method on all datasets. meanAUC is the unweighted average of AUC scores over all proteins in a dataset. In all cases the naive performance is meanAUC of 0.5.

While all interpretability methods achieve positive signal on most datasets, they provide largely non-specific results, with LDL and LGA achieving meanAUC on the order of 0.6 for most datasets table 3. Qualitatively, readouts of interpretability scores show very noisy relevance predictions, with a slight bias towards positively annotated sites fig. 3 B. Interpretability methods do show qualitatively similar prediction patterns, and output highly correlated residue rankings fig. 3 B, indicating extraction of similar data from the base classifier.

**Fig. 3:**
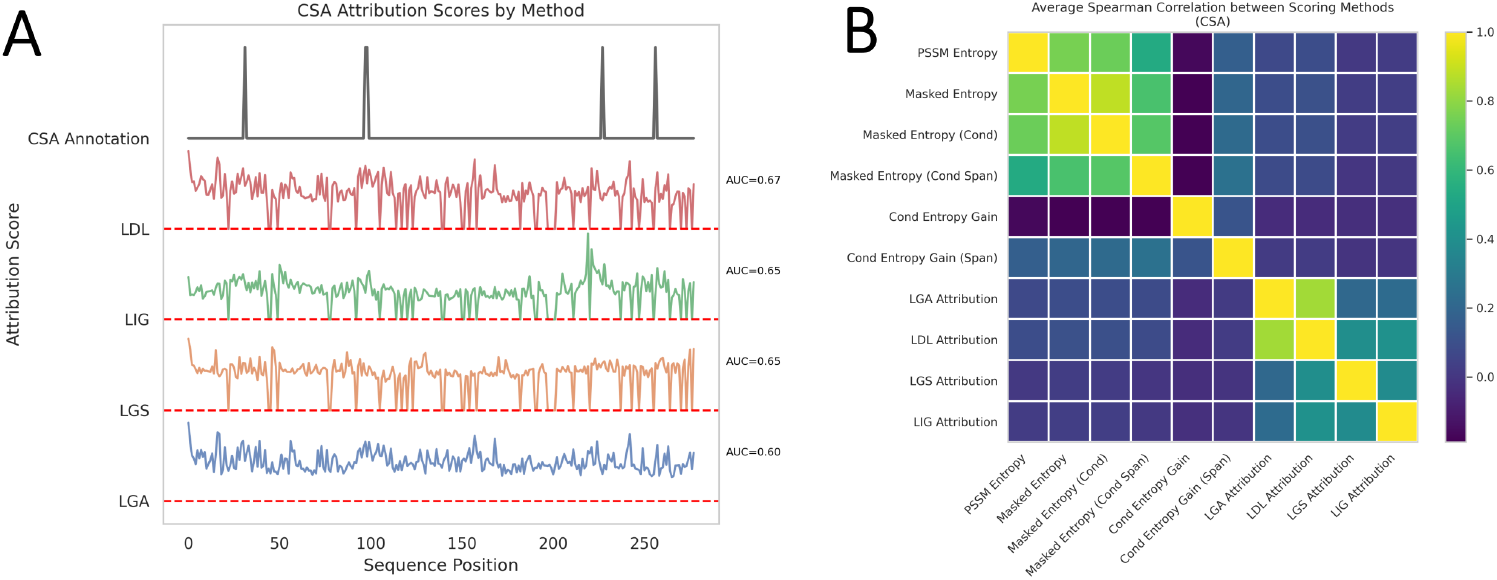
A) Case study of interpretability methods predicting residue importances. Layer attribution methods applied to UniProt O31168 protein to estimate contribution of residues to a positive prediction of chloride peroxidase activity. B) Averaged Spearman’s rho coefficients of residue rankings from different methods on the CSA dataset.

**Fig. 4:**
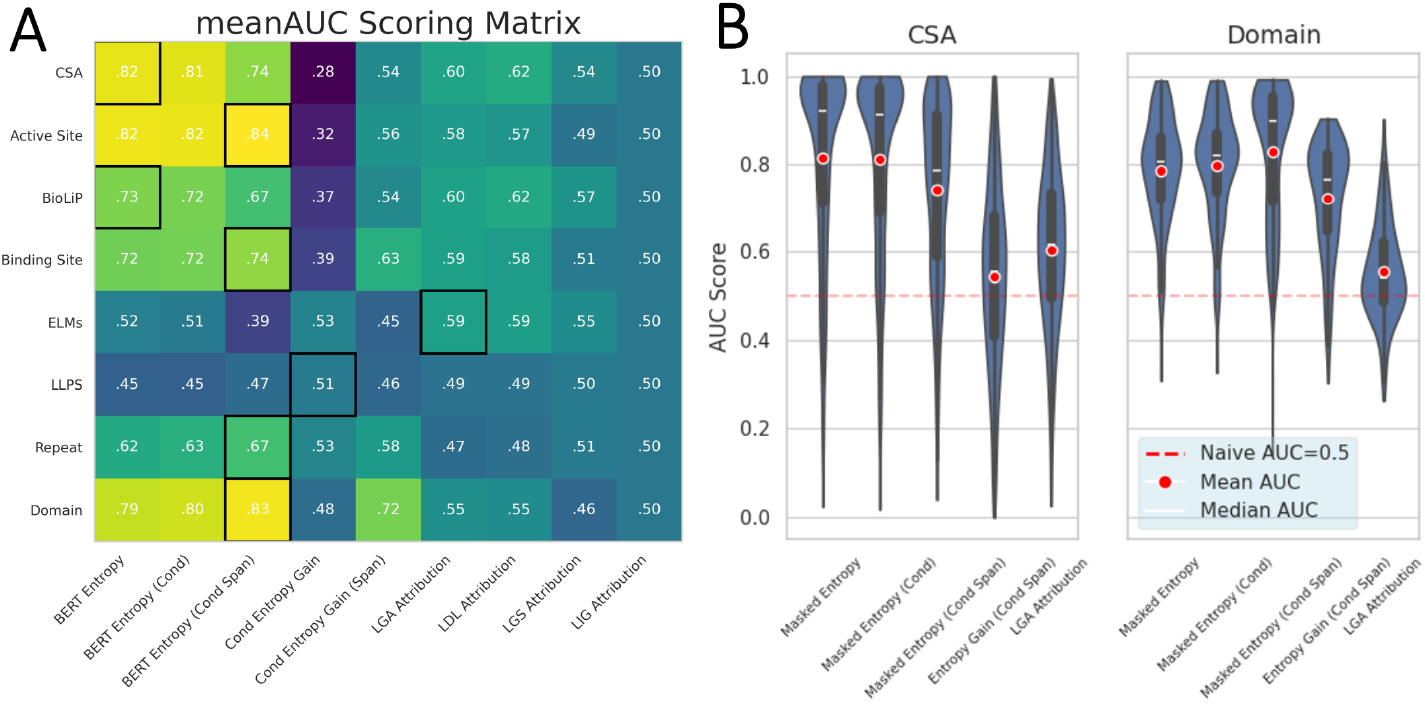
A) All-dataset comparison of Masked Entropy, Function-Conditioned Entropy, and interpretability methods on retrieval of functionally important residues, as evaluated by meanAUC. Highest performing model outlined in black for each dataset. B) Violin-plots of the distribution of per-protein AUC scores. Mean and median meanAUC plotted for each model in red dot and white dash respectively.

### 3.3. Retrieval Comparison on All Datasets

In our final comparison, we evaluate both ESM-C derived entropy estimation methods and classifier interpretability methods on each of our constructed datasets.

Because of its applicability to all datasets, at all annotation specificity levels, we use meanAUC scores for comparison between methods. While Top30 metrics show similar results, MRR metrics tend to favor demasking entropy models with higher performance on outlier residues such as active site amino acids, as shown in Figure S1.

#### 3.3.1. Naive Entropy Methods Dominate for Highly Specific Retrieval

During our full comparison, we find that for both the CSA and BioLiP datasets it is most effective to score residues by entropy estimates, without regard for function. We hypothesize that for extremely important residues within a protein, such as active sites or critical residues within binding sites, conservation is sufficiently high that functional information is redundant. In these cases, direct masked entropy estimates dominate. However, they show much poorer performance for potentially low entropy regions, in particular for repeat and domain regions extracted from the InterPro dataset.

#### 3.3.2. Functional Information Improves Retrieval For Functional Regions

In most cases, function conditioned entropy performs similarly to entropy derived from our pre-trained model. We hypothesize that under the default 15% masking scheme used for residue distribution estimation functional information is largely redundant. This is supported by the performance of conditional entropy gain as a retrieval metric, in which gain under default masking correlates negatively with entropy based predictions fig. 3, and worse than random on most tasks.

With the introduction of context-span masking, we find that our model gains significantly more information from protein function. The span-masked model achieves the highest performance out of all models on all InterPro derived datasets section 3.3. In addition, entropy-gain derived from the span-masked and span-mask baseline models achieves significant positive signal on all but the ELMS and LLPS datasets, and outperforms baseline interpretability methods section 3.3.

Qualitatively, such as in fig. 5, we see that the most significant areas of entropy gain are in functionally relevant regions. Entropy-gain may be thought of as an additional, functionally determined, signal, which may be added to the baseline entropy prediction to produce a more discriminative prediction.

**Fig. 5:**
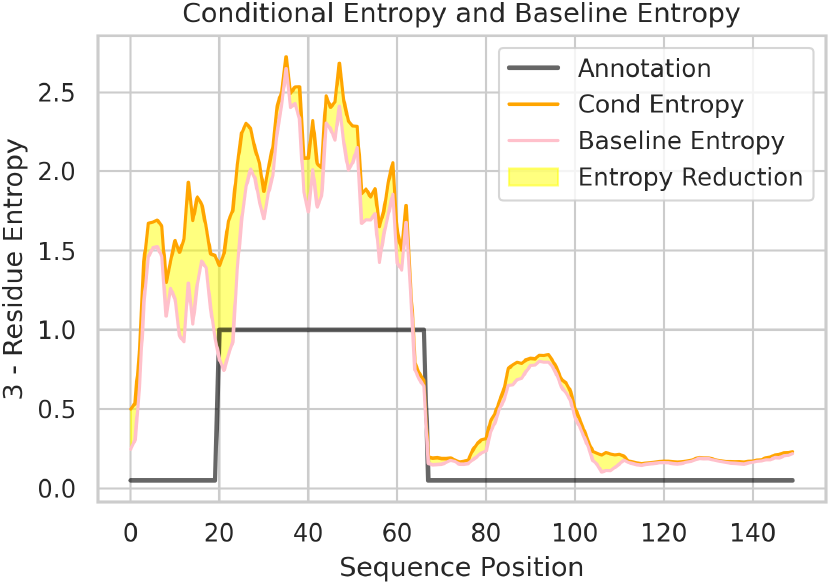
Casestudy of the effect of functional information on residue entropy. A 3 − *h*_*θ*_(*r*_*i*_) score plotted for both function conditioned and baseline residue entropy estimates. Functional information lowers model entropy on function relevant regions, increasing retrieval rate.

One of the costs of the span-masking objective is diminished scoring on the highest specificity datasets such as CSA. Missing information, and missing local context from the masking pattern, increases overall model uncertainty on highly specific residues. In almost all cases, interpretability based methods derived from a classifier perform poorly for residue retrieval table 3.

The exception is for the ELMS dataset, in which layer gradient activation and layer deep lift both achieve significant signal, and convincingly outperform all other methods. Notably, the ELMS dataset is composed of relatively high entropy residues, which may not correspond to truly low entropy regions even for a perfect model with access to functional information. This may represent an adversarial example for direct sequence conservation based methods.

We also find that all datasets perform with close to random performance on the LLPS dataset, in which disordered regions represent a challenging target for both classifiers and generative models. High performance on the LLPS dataset may require explicit treatment of disordered regions, or more powerful models of the sequence distribution.

## 4. Discussion

Determining the functional importance of protein residues is a critical, but at the same time fundamentally ill-posed, problem. Proteins may have multiple functions Jeffery 2018, critical structural elements Ismi, Pulungan, and Afiahayati 2022, or dozens of other conserved properties that may be so entangled with function that they are inseparable.

In agreement with previous research into sequence conservation Capra and Singh 2007, we find that residue entropy derived from PSSM estimates shows extremely high correlation with residue functional relevance, especially in the case of the most highly conserved locations such as catalytic sites. On other datasets, with the notable exception of ELMS motif sequences, they show similarly strong performance, despite using no functional information whatsoever in their predictions.

Our results indicate that the greater availability, and higher information density, of sequence datasets allows sequence based models to achieve high performance on residue retrieval tasks. The high performance of MLM entropy supports a long-running trend in bioinformatics research Zuckerkandl and Pauling 1965, Ashkenazy, Erez, et al. 2010, Capra and Singh 2007, in residue entropy is a very strong heuristic indicator for functional relevance. Across all datasets and metrics, we observe better retrieval from MLM sequence models than MSA alignments, with the ESM-C model showing better generalization than direct alignment of homologous sequences. The extremely high correlation of ESM-C and MSA estimates of residue distributions indicates that both are estimating similar protein sequence properties from similar data, but that the higher flexibility of direct generative modeling eventually produces stronger results.

In the case of regions which may not be uniformly conserved such as domains, repeat regions, or less specific active or binding sites, functional information becomes more important for specific retrieval. For these cases, we demonstrate that knowledge of a protein’s intended function can augment generative models. While function information shows little benefit for models trained on a typical 15% masking objective, it significantly influences models trained on the more challenging context-span masking objective. We confirm that these function-conditioned models have the highest retrieval performance for broader annotation types, and through a comparison to baseline models we verify that the reduction in residue entropy from this functional knowledge constitutes a signal for relevance that is independent of sequence conservation.

A critical limitation for all methods is their failure in cases of high sequence disorder, as is the case for ELMS motifs and intrinsically disordered domains from the LLPS dataset. Although these regions are experimentally confirmed to be critical for protein function Mészáros et al. 2020, Kumar et al. 2019, they do not show significantly low entropy. All methods perform comparably to random chance on the LLPS dataset, while only interpretability methods achieve significant signal on the ELMS dataset.

A fully-generative model, trained from scratch to reconstruct function and sequence simultaneously, would likely show stronger performance in both cases, allowing for inference of both region and function simultaneously. Modified generative architectures such as auto-regressive generative models Shin et al. 2021, or masked diffusion models Hallee et al. 2025 would allow for the calculation of entropy jointly over entire regions, potentially addressing current difficulties in modeling difficult targets such as the LLPS dataset.

## 5. Conclusion

In this work, we formalize unsupervised functional residue annotation through a benchmark suite of eight datasets. We demonstrate that while entropy from pre-trained language models is a strong baseline for conserved residue identification, it may be outperformed by a function-conditioned generative model. When trained with a novel context-span masking objective, this model excels at retrieving broader functional regions like domains and repeats. We also introduce entropy gain, calculated as 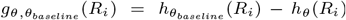, as a method to specifically isolate the impact of functional information on a residue’s predicted distribution. This gain metric provides a distinct signal of functional relevance, independent of sequence conservation.

The overall pipeline offers a purely unsupervised framework for identifying functionally important domains, motifs, or sites within proteins of interest. In the future we hope to see successes in function-conditioned modeling applied directly to the guidance of residue-level protein experiments, or even direct protein engineering through residue optimization.

## 7. Author contributions statement

A.D. and M.M. conceived the experiments, A.D. and S.M. conducted the experiments and analyzed the results. A.D., S.M. and A.T. assembled annotation datasets. A.D. and M.M. wrote and reviewed the manuscript.

## 8. Acknowledgments

We thank Salmonn Talebi, Ehsaneddin Asgari, and other members of the Molecular Cell Biomechanics Laboratory for the suggestions and comments they provided throughout this research.

