## Supplementary Information for "Semi-supervised Retrieval of Functional Residues Through the Integration of Protein Language Models and Gene Ontology Data"

### 1. Training Methodology

The following outlines the computational methodology used to train the models described in the main paper. Our work centered on two main modeling approaches: (1) a Gene Ontology (GO) classifier, which was fine-tuned to serve as a basis for evaluating classifier interpretability methods, and (2) a function-conditioned generative model, developed to directly integrate functional information into residue-level predictions. All models were implemented using the PyTorch and PyTorch Lightning frameworks.

#### 1.1. Gene Ontology (GO) Classifier for Interpretability

To establish a baseline and evaluate the efficacy of feature attribution techniques, we fine-tuned a protein language model for multi-label GO term classification.

The classifier was built upon the pre-trained 600M ESM-Cambrian (ESM-C) protein language model. For a given input sequence, the final-layer residue embeddings from the ESM-C encoder were subjected to mean pooling to produce a single, fixed-size vector representing the entire protein. This sequence-level representation was then fed into a linear classification head to produce logits for each GO term in our dataset.

#### 1.2. Training and Optimization

We adopted a two-phase fine-tuning strategy on our curated SwissProt-GOA dataset. 1. Phase 1 (Head Training): The parameters of the ESM-C base model were frozen, and only the final linear classification head was trained. This phase used a higher learning rate to quickly adapt the new layer to the task. 2. Phase 2 (Full Model Fine-tuning): The entire model, including the ESM-C encoder, was unfrozen and trained for five epochs. We used the AdamW optimizer with a learning rate that was annealed from $1 \times 10^{-5}$ down to $1 \times 10^{-6}$.

The model was trained by minimizing the Binary Cross-Entropy (BCE) loss, and its performance was monitored using the micro F1-score on a held-out validation set.

#### 1.3. Function-Conditioned Generative Model

This model was developed to guide the generative process of a protein language model with functional annotations, with the goal of improving the identification of functionally relevant residues.

##### 1.3.1. Model Architecture

The architecture augments the pre-trained ESM-C model. A learnable GO term embedding matrix, $E_{GO} \in \mathbb{R}^{|\mathcal{G}| \times d_{emb}}$, was introduced, where $|\mathcal{G}|$ is the total number of GO terms and $d_{emb}$ is the model's embedding dimension. For a protein annotated with a set of GO terms represented by a multi-hot vector $y$, an aggregate function vector $v_{func}$ was computed by summing the corresponding embedding vectors:

$$v_{func} = \sum_{j=1}^{|\mathcal{G}|} y_j E_{GO}^{(j)}$$

This vector was then passed through a linear projection and added to each residue embedding at the input layer of the ESM-C transformer. This allows the model to condition its sequence predictions on the global functional context provided.

#### 1.3.2. Training Objectives and Masking Strategies

The model's parameters (the ESM-C base and the new function embeddings) were jointly fine-tuned using a masked language modeling (MLM) objective. We experimented with two distinct masking schemes:

In the default training, following the original BERT methodology, 15% of the residues in each sequence were randomly selected and replaced with a '[MASK]' token. In Context-Span Masking, the objective is made more challenging to increase the relevance of functional information. For each sequence, a random, contiguous 100-residue "focus region" was defined. All residues outside this region were masked and excluded from the loss calculation. Within the focus region, contiguous spans of 2 to 5 residues were masked until 15% of the total sequence residues were masked. The model was trained to predict the original amino acids for these masked spans.

#### 1.3.3. Training and Optimization

The function-conditioned model was fine-tuned for five epochs using the AdamW optimizer. The learning rate was annealed from an initial value of $1\times10^{-5}$ to a final value of $1\times10^{-6}$. The training loss was the standard cross-entropy loss calculated over the masked positions. A baseline model (standard ESM-C without functional conditioning) was also fine-tuned under identical conditions for comparison.

### 2. Derivation of Residue Importance Scores

To evaluate the models, we derived a per-residue importance score from each.

#### 2.1. Positional Entropy

For a given sequence, a Position-Specific Scoring Matrix (PSSM) was estimated by repeatedly masking each position and averaging the model's output probability distributions for that position. The final importance score was the Shannon entropy of this predicted distribution, where lower entropy signifies higher conservation and thus greater importance.

From Function-Conditioned Models (Entropy Gain): To isolate the contribution of the functional information, we calculated the entropy gain. This metric is defined as the reduction in positional entropy when functional conditioning is applied: $g_{\theta,\theta_{baseline}}(R_i) = h_{\theta_{baseline}}(R_i) - h_\theta(R_i)$. A positive gain indicates that the functional context made the model more certain about a residue's identity.

#### 2.2. Masking Strategies

Estimating PSSMs from our generative models requires us to repeatedly mask and demask protein residues to predict their conservation. We evaluate strategies of both randomly masking a percentage of residues for each demasking iteration and iterating through individual residues and masking only them. We also evaluate a span-masking strategy in which as sliding window of masked residues is run across the protein.

For percentage masking trials, residues may be evaluated any number of times. We test coverage levels of 1, 6, and 12, to evaluate the effect of repeated trials.

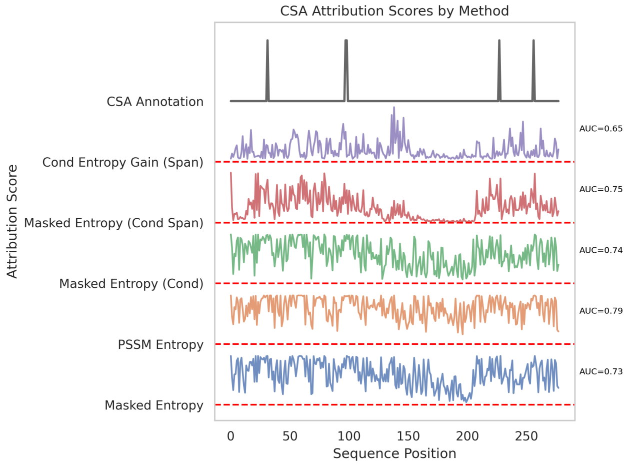


Fig. 1: Case study of residue entropy based methods predicting residue importances of Uniprot protein O31168. Residue number plotted against normalized importance scores corresponding to negative entropy in the case of entropy based methods, or reduction in entropy in the case of entropy gain methods.

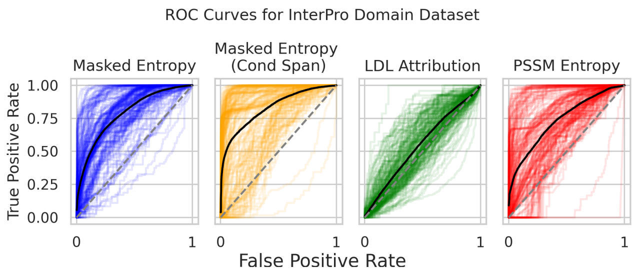


Fig. 2: ROC curves for domain retrieval from the InterPro domain dataset on each dataset protein, plotted for the masked entropy, masked entropy with context-span masking and functional conditioning, LDL attribution, and PSSM entropy.

#### 2.3. Interpretability

From the GO Classifier (Interpretability Methods): We applied four feature attribution methods to the fine-tuned GO classifier: Layer-wise Integrated Gradients (LIG), Layer-wise Gradient X Activation (LGA), Layer-wise DeepLIFT (LDA), and Layer-wise GradientShap (LGS). For each method, the L2 norm of the attribution values for a residue's input embedding was taken as its final importance score.

**Table 1.** Performance metrics for the CSA dataset. MRR, AUC, BAUC, and Top-30 metrics reported. BAUC represents AUC values of ROC curve over all residues.

| Method | Perc | Coverage | MRR | AUC | BAUC | Top-30 Score |
|---|---|---|---|---|---|---|
| Percentage Masking | 0.05 | 1 | 0.1336 | 0.7980 | 0.7804 | 0.5484 |
| Percentage Masking | 0.05 | 6 | 0.4100 | 0.8147 | 0.7954 | 0.5940 |
| Percentage Masking | 0.05 | 12 | 0.4091 | 0.8153 | 0.7956 | 0.5961 |
| Percentage Masking | 0.15 | 1 | 0.0401 | 0.7308 | 0.7173 | 0.1160 |
| Percentage Masking | 0.15 | 6 | 0.4154 | 0.8160 | 0.7954 | 0.5946 |
| Percentage Masking | 0.15 | 12 | 0.4130 | 0.8158 | 0.7952 | 0.5970 |
| Percentage Masking | 0.3 | 1 | 0.0415 | 0.7315 | 0.7162 | 0.1289 |
| Percentage Masking | 0.3 | 6 | 0.4112 | 0.8145 | 0.7917 | 0.5913 |
| Percentage Masking | 0.3 | 12 | 0.4142 | 0.8167 | 0.7932 | 0.5966 |
| Indiv Masking | - | - | 0.4088 | 0.8147 | 0.7953 | 0.5950 |
| Neighborhood 5 Masking | - | - | 0.4040 | 0.7999 | 0.7749 | 0.5467 |
| Naive | - | - | 0.0589 | 0.5006 | 0.5005 | 0.0909 |

**Table 2.** Performance metrics for the IP domain dataset. MRR, AUC, BAUC, and Top-30 metrics reported. BAUC represents AUC values of ROC curve over all residues.

| Method | Perc | Coverage | MRR | AUC | BAUC | Top-30 Score |
|---|---|---|---|---|---|---|
| Percentage Masking | 0.05 | 1 | 0.2651 | 0.7575 | 0.7525 | 0.5256 |
| Percentage Masking | 0.05 | 6 | 0.7827 | 0.7801 | 0.7730 | 0.6879 |
| Percentage Masking | 0.05 | 12 | 0.7826 | 0.7801 | 0.7732 | 0.6899 |
| Percentage Masking | 0.15 | 1 | 0.1552 | 0.6665 | 0.6742 | 0.1602 |
| Percentage Masking | 0.15 | 12 | 0.7834 | 0.7865 | 0.7789 | 0.7008 |
| Percentage Masking | 0.15 | 6 | 0.7848 | 0.7858 | 0.7783 | 0.6974 |
| Percentage Masking | 0.3 | 1 | 0.1572 | 0.6759 | 0.6816 | 0.1631 |
| Percentage Masking | 0.3 | 6 | 0.7896 | 0.7933 | 0.7856 | 0.7101 |
| Percentage Masking | 0.3 | 12 | 0.7875 | 0.7944 | 0.7865 | 0.7084 |
| Indiv Masking | - | - | 0.7753 | 0.7771 | 0.7699 | 0.6837 |
| Neighborhood 5 Masking | - | - | 0.8185 | 0.8099 | 0.7990 | 0.7417 |
| Naive | - | - | 0.5026 | 0.4959 | 0.5004 | 0.3237 |

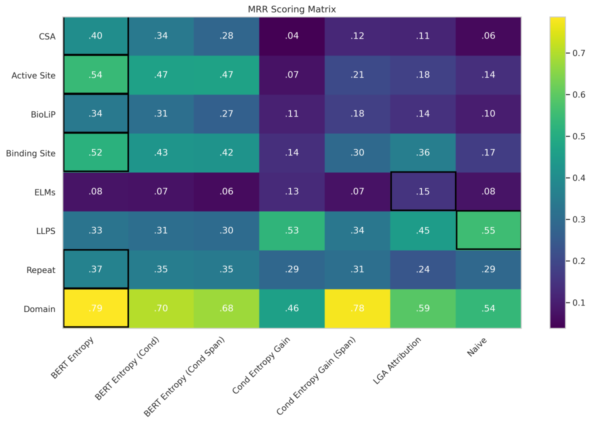


Fig. 3: A) All-dataset comparison of Masked Entropy, Function-Conditioned Entropy, and interpretability methods on retrieval of functionally important residues, as evaluated by Mean Reciprocal Rank.

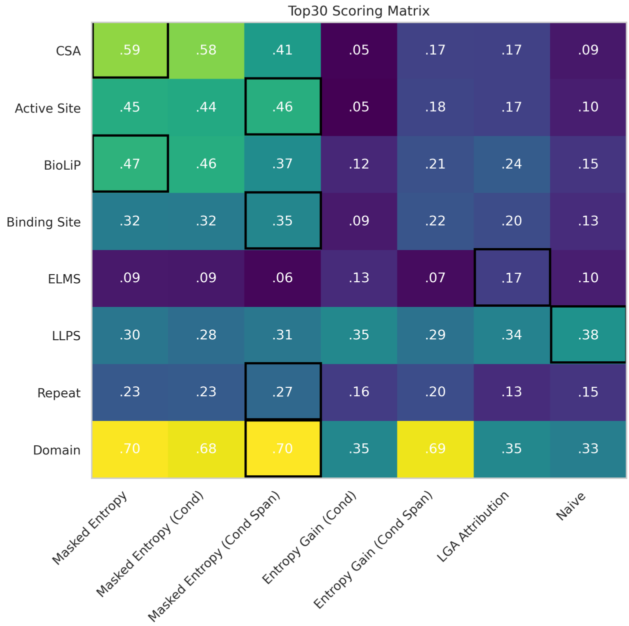


Fig. 4: A) All-dataset comparison of Masked Entropy, Function-Conditioned Entropy, and interpretability methods on retrieval of functionally important residues, as evaluated by mean of Top30 metric across dataset.

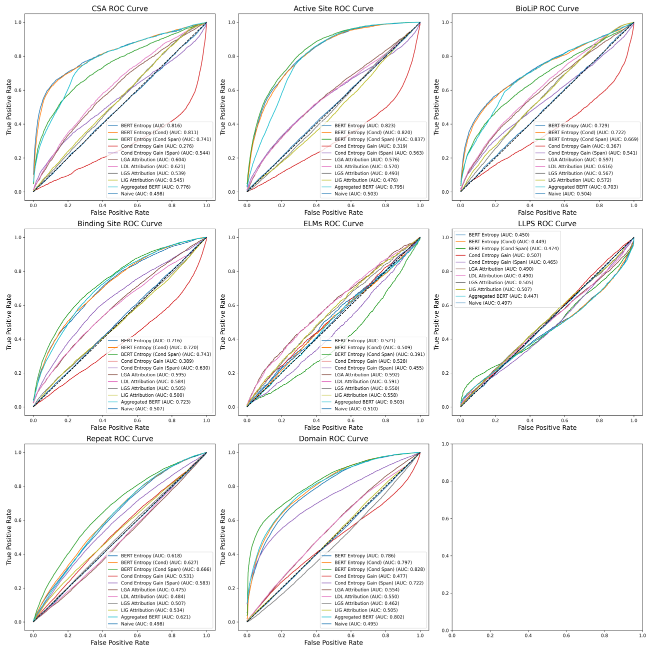


Fig. 5: Plots of meanROC curves for all models on all datasets.

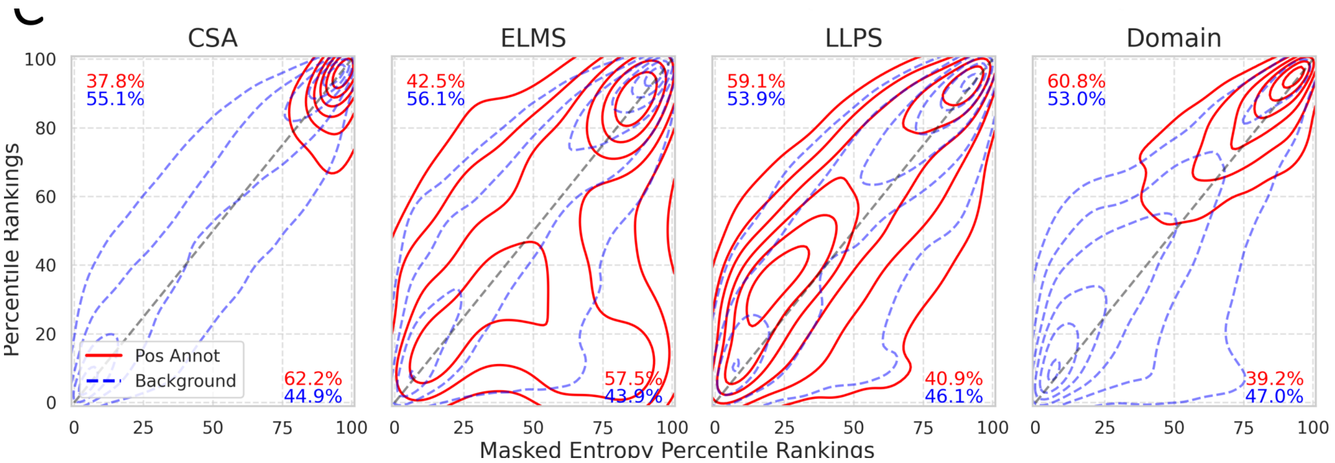


Fig. 6: Topographic maps of percentile scoring by ESM-C masked entropy vs. ESM-C fine-tuned with a functional context on a context-span masking loss. For each sequence in our dataset, percentile ranking of each residue as scored by Masked Entropy and Masked Entropy with functional conditioning and context-span masking are calculate. We plot the density of residues from all sequences, inversely weighted by sequence length, on the space of pairs of percentile importance assignments from the two models. Density plots for annotated and baseline residues plotted separately, with percentage of residues for which entropy rank is higher than span-context entropy rank labeled in lower-right, and inverse labeled in upper-left.
